# Learning the Malignant Transformation of Lung Adenocarcinoma Cells Using Explainable AI

**DOI:** 10.64898/2026.11.25.769087

**Authors:** Doreen Klingler, Linda Bergmayr, Konstanze Schleich, Mina Jamshidi Idaji, Edgar Chimal Calderon, Liliana H Mochmann, Yauheniya Zhdanovich, Fabian Engelhardt-Schott, Julius Keyl, Andreas Mock, Klaus-Robert Müller, Frederick Klauschen, Philipp Jurmeister, Philipp Keyl

## Abstract

Tumor evolution in lung adenocarcinoma (LUAD) drives drug resistance and disease progression, yet the cellular transitions underlying the malignant transformation of cells remain poorly understood. Comparing this progression of cancer cells across patients is challenging owing to substantial inter-patient cell heterogeneity. Here, based on single-nucleus RNA sequencing (snRNA-seq) data of 227 patients with LUAD including 422,727 tumor cells, we conducted a comprehensive investigation of tumor cell progression and developed Global Tumor Time (GTT), a continuous measure of tumor cell progression across patients. GTT provided a scalable and biologically meaningful metric of tumor progression that aligned with clinicopathological, transcriptomic and spatial characteristics of malignancy and demonstrated robust prognostic performance (C-index = 0.60), which was further validated in two external datasets comprising snRNA-seq (C-index = 0.66) and bulk RNA sequencing (C-index = 0.61). Using explainable AI to investigate the molecular correlates of GTT, we identified two distinct transcriptional programs associated with early and late progression states. These findings establish a scalable and interpretable framework for reconstructing malignant cell transformation across patients and allow for investigating the biology of tumor progression at single-cell resolution.

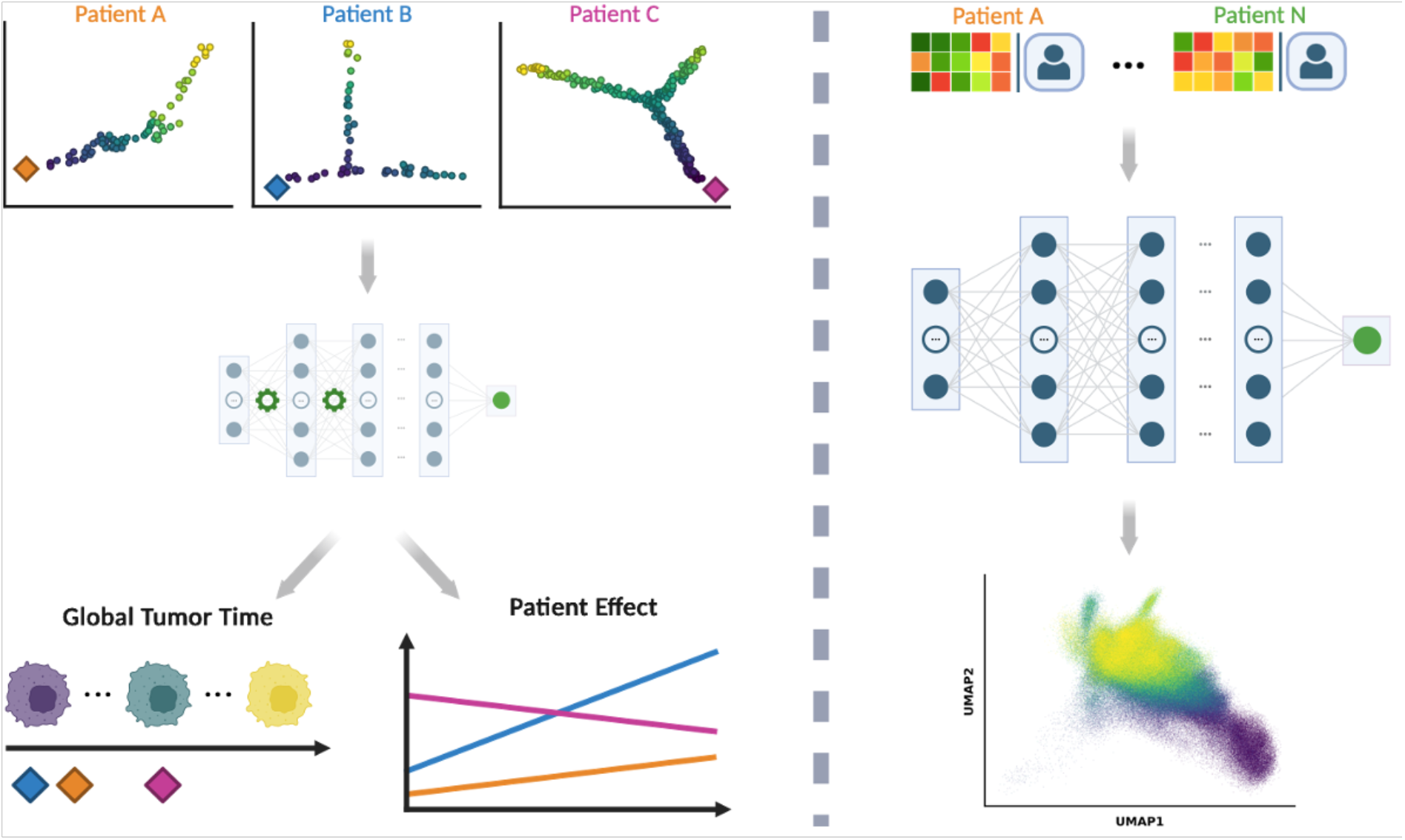

## Introduction

Lung adenocarcinoma (LUAD), the most common subtype of lung cancer, is characterized by profound intra- and inter-tumor heterogeneity that limits durable therapeutic response and contributes to disease progression^1,2^. This heterogeneity arises from ongoing clonal evolution, a multi-step process through which subpopulations of tumor cells progressively acquire enhanced malignant traits^3,4^. Understanding the progression of individual tumor cells is therefore essential for quantifying tumor heterogeneity, predicting therapy resistance, and guiding personalized treatment strategies^5–7^.

Single-cell RNA sequencing (scRNA-seq) provides the opportunity to study tumor evolution at cellular resolution by capturing transcriptional states of thousands of individual cells^8,9^. Pseudotime analysis has emerged as a computational approach that infers a continuous trajectory from scRNA-seq data and assigns each cell a relative position along this trajectory with respect to a predefined root cell. This ordering approximates cellular progression during processes such as differentiation or malignant transformation^10^.

As most methods infer pseudotime values within individual samples, direct comparisons of pseudotimes across multiple patients or conditions remains challenging^11^. Previous pseudotime analyses of cancer cell progression in LUAD have therefore either relied on small scRNA-seq cohorts with prior batch-correction^12–16^ or on bulk transcriptomic data using methods such as PhenoPath^17–19^. However, enforcing a shared trajectory through batch-correction may obscure inter-patient variability. Additionally, many methods that attempt to define common trajectories of cell (de-)differentiation lack scalability to large scRNA-seq cohorts and assume linear relationships between gene expression and pseudotime^11,17^, limiting their ability to capture the complex dynamics of tumor evolution.

In this study, we conducted a comprehensive analysis of tumor cells across 227 LUAD patients to characterize tumor cell progression on a common scale across individuals. We developed a neural network framework, GTTimer, that extracts a global tumor time (GTT) directly from single-cell gene expression by decomposing each cell’s pseudotime into GTT and patient-specific effects. We demonstrate that GTT is strongly associated with established clinicopathological, transcriptional, and spatial markers of tumor progression. Moreover, mean GTT per patient is predictive for overall survival, remains robust across two independent external validation cohorts, and outperforms existing approaches in predictive performance and scalability. Using the xAI method layer-wise relevance propagation (LRP)20, we further provide an interpretable framework that identifies genes driving tumor progression.

This work offers a biologically grounded framework for understanding how tumor cells evolve from normal epithelium and progress toward advanced malignant states. The tumor progression trajectory inferred by GTTimer provides a foundation for dissecting conserved and patient-specific programs of LUAD evolution, with potential implications for prognostic assessment and therapeutic decision-making.

## Results

### A deep learning model for Global Tumor Time prediction

To investigate the dedifferentiation and progression of cancer cells across patients, we leveraged the LuCaS+ atlas^20,21^, comprising single nucleus RNA sequencing (snRNA-seq) data from 227 patients with primary LUAD spanning UICC stages I-IV, and filtered for epithelial malignant and non-malignant cells. We computed diffusion pseudotime^22^ (DPT) independently for each patient, using malignant cells and type II pneumocytes and defining the average non-malignant epithelial cell within each sample as the root cell.

Since pseudotime provides only a local and inherently relative measure of progression, we first investigated the use of gene-wise linear mixed-effects to uncover the overall association (fixed effects) between the respective gene and the transcriptomic distance between malignant and normal epithelial cells across all patients. Meanwhile, the patient-specific random effects accounted for the patient-specific diffusion pseudotime scale between zero and one. The fixed-effect estimates, restricted to the top 10% of genes with the best model performance (lowest RMSE), showed a significant correlation with disease progression, were reproducible in an external dataset, and showed prognostic relevance in LUAD patients in the TCGA dataset (Supplementary Figure S1). However, while the selected genes provided robust markers of tumor progression across datasets, they offer only univariate, linear estimates of tumor cell trajectories, highlighting the need for a nonlinear, unified measure of cellular progression based on their joint expression.

To this end, we developed GTTimer, a multivariate neural network-based framework that extends the mixed-effects modelling approach and infers a unified temporal score, Global Tumor Time (GTT), from the expression of 5,000 genes. Using DPT as a supervisory signal, GTTimer was trained on 192 patients and validated on 35 patients to derive GTT by explicitly modeling patient-specific effects, yielding a common progression axis that enables direct comparison of tumor cells across patients.

Mean GTT per patient correlated with important clinical tumor characteristics indicating tumor progression and dedifferentiation of cancer cells, such as staging (Spearman’s rho = 0.21, p = 0.002), grading (Spearman’s rho = 0.48, p < 0.001) and the presence of lymph node metastases (Spearman’s rho = 0.17, p = 0.011; Fig. 1B), indicating that GTT reflects clinically relevant disease severity. On a molecular level, stratification of individual tumor cells by GTT revealed a progressive shift of cellular proportions from a predominantly resting state toward a proliferative state with increasing GTT (Fig. 1C), indicating enhanced cell cycle activity in tumor cells with higher GTT. Hypoxia scores showed a similar increase with higher GTT (Spearman’s rho = 0.30, p < 0.001; Fig. 2D). Beyond functional state changes, higher GTT was associated with higher CNV scores (Spearman’s rho = 0.54, p < 0.001; Fig. 1E), and greater inferred cellular potency (Spearman’s rho = 0.58, p < 0.001; Fig. 1F), suggesting increased genomic instability^23^ and a transition towards a less differentiated, more stem-like phenotype.

**Figure 1:**
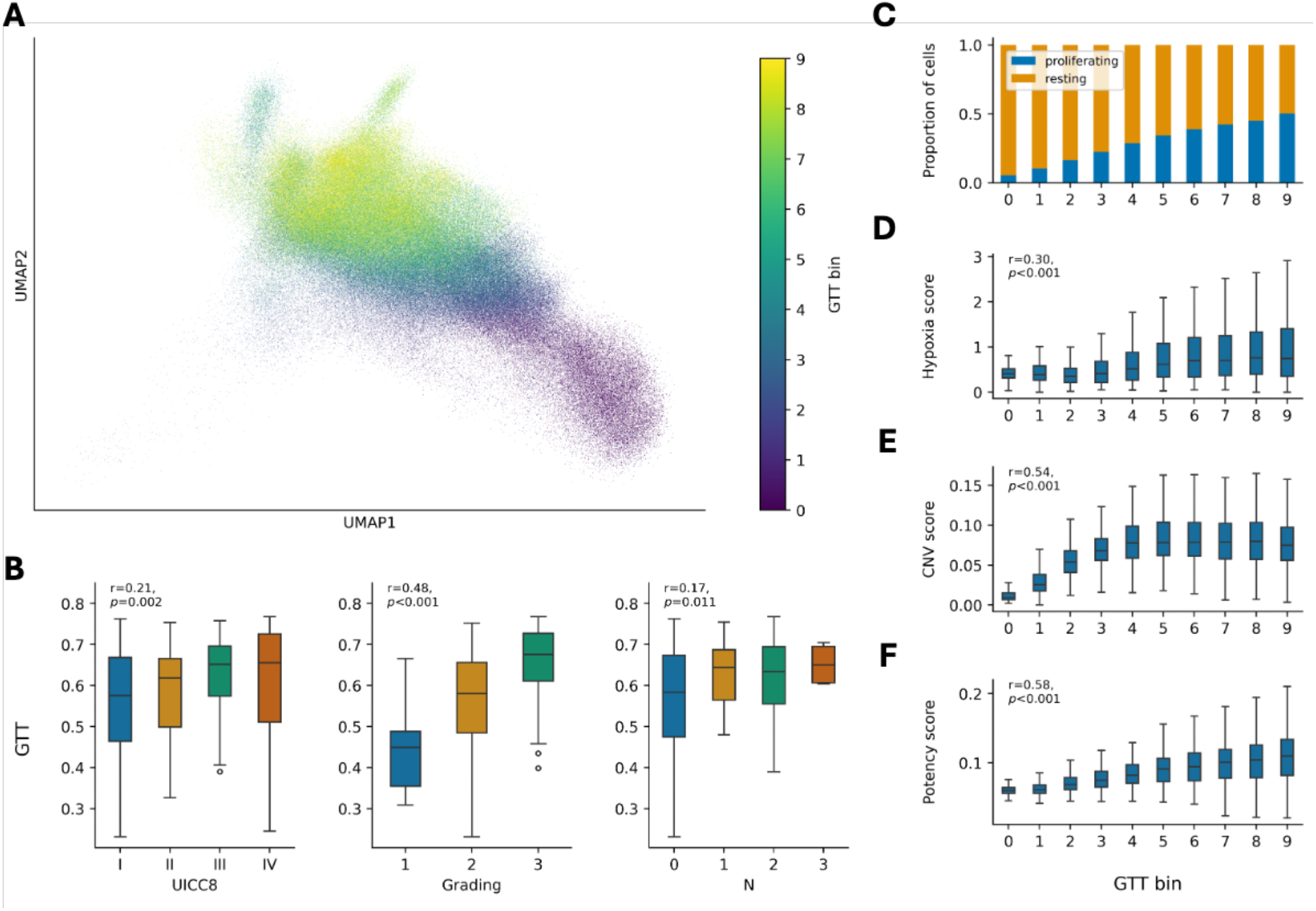
GTT is associated with clinicopathological characteristics and tumor-related transcriptional programs. (A) UMAP embedding of the analyzed cells. Colors indicate GTT bins (0-9). (B) Mean GTT per patient across clinicopathological features, including UICC stage (8th edition), tumor grade, and nodal status (N). (C–F) Relationships between binned GTT and (C) cell-cycle activity, based on cell cycle phase assignment using the expression of S- and G2/M-phase genes; (D) hypoxia, defined as the mean expression of Tardon21 signature genes; (E) CNV scores inferred by inferCNV (CNV score is defined as the fraction of genes with expression deviating from the normal baseline (1.0) by more than ±5%); and (F) cellular potency score inferred by CytoTRACE2 (higher scores correspond to less differentiated, more stem-like cell states). Spearman correlation coefficients (r) and corresponding p-values (p) are indicated in the panels.

### Spatial distribution of GTT

Histopathological assessment is the current standard for distinguishing normal epithelial cells from dysplastic or invasive tumor cells and for grading tumors over the course of disease progression. We therefore investigated whether the measure of tumor cell progression captured by GTT also manifests in histomorphological characteristics using 152 matched Xenium spatial transcriptomics samples with GTT transferred from snRNA-seq, and H&E-stained histopathology data. GTT showed a positive correlation with cell (Pearson’s r = 0.24, p < 0.001) and nucleus size (Pearson’s r = 0.18, p < 0.001), consistent with continuous alteration of cell morphology with increasing GTT (Fig. 2A, B, C). In contrast, GTT was negatively correlated with the distance to neighboring tumor cells (Pearson’s r = −0.17, p < 0.001), consistent with higher tumor cell density as observed in solid - by definition high-grade - growth patterns of LUAD (Fig. 2D). This was in line with an analysis of the neighborhood composition, defined as all cells within a 30 μm radius of each tumor cell (Fig. 2E). Tumor cells with high GTT were positioned predominantly in tumor cell-enriched microenvironments (Fig. 2E), while the density of immune cells declined. Together, these findings indicate that advanced tumor cells are characterized by increased spatial aggregation and reduced immune presence, consistent with an increasingly immune-evasive microenvironment.

**Figure 2:**
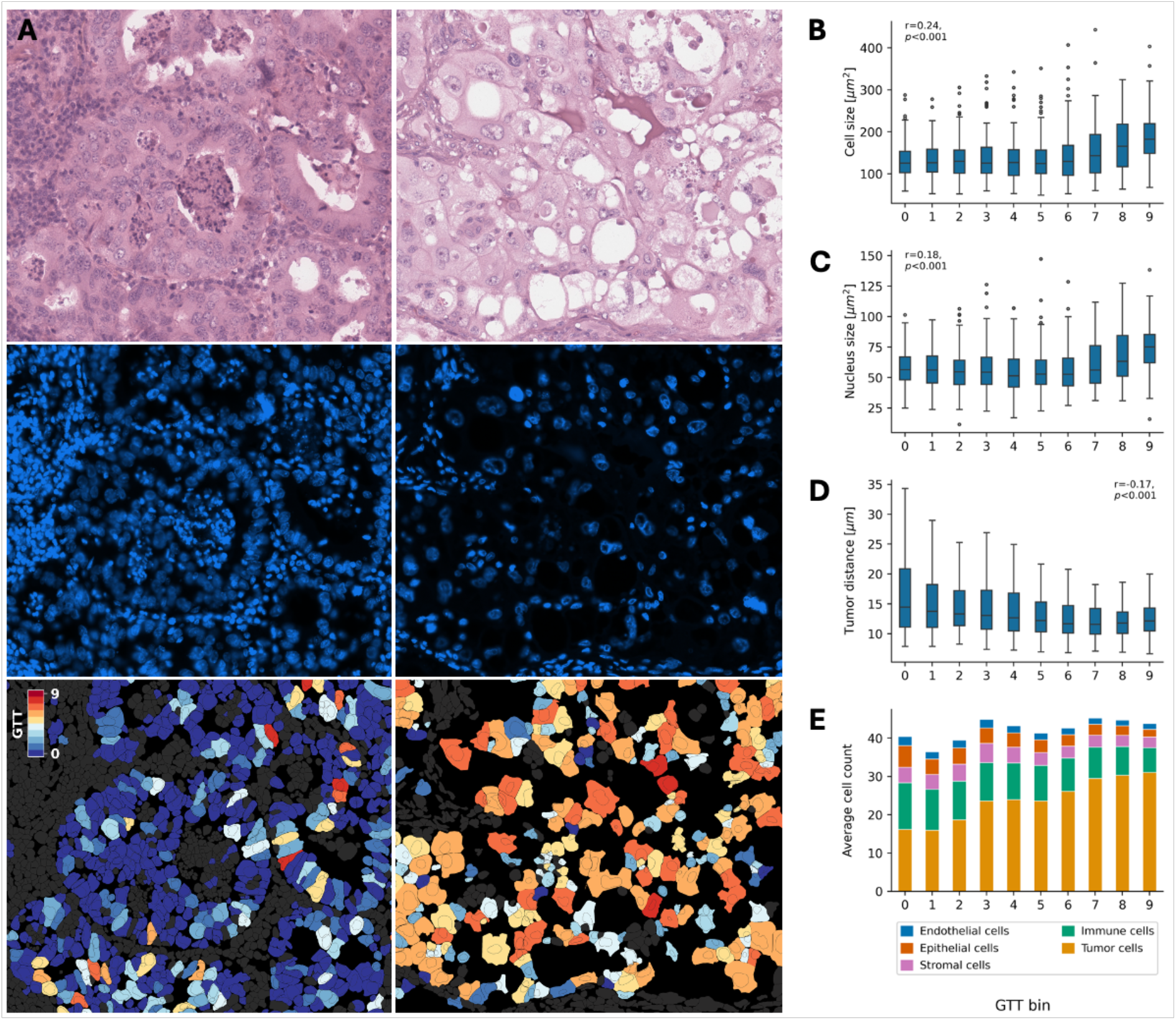
Spatial transcriptomics demonstrates that GTT is associated with increased tumor cell and nucleus size and microenvironmental features. (A) Representative hematoxylin and eosin (H&E) images (top), corresponding DAPI-stained nuclei (middle), and spatial distribution of GTT (bottom) for regions with high (left) and low (right) GTT. High-GTT regions exhibited heterogeneous nuclear sizes, abundant cytoplasm, and prominent nucleoli, whereas low-GTT regions showed more uniform nuclear morphology and inflammatory infiltration at the tissue borders. (B) Correlation between GTT and cell size. (C) Correlation between GTT and nucleus size. (D) Correlation between GTT and distance to the nearest tumor cell. (E) Composition of cell types within the tumor neighborhood (30 µm radius around each tumor cell). Pearson correlation coefficients (r) and corresponding p-values (p) are indicated where applicable.

### Association of GTT with remodeling of the immune landscape

To determine whether the observed reduction in immune cell abundance with increasing GTT was accompanied by compositional changes, we next analyzed immune cell subtype proportions (Fig. 3). Higher GTT was associated with increased proportions of plasma cells, accompanied by reduced dendritic cells (Pearson’s r = -0.94, FDR q-value < 0.001), indicating a shift in local immune composition.

Within the T cell compartment, proliferating (cycling) cytotoxic (Pearson’s r = 0.96, FDR q-value < 0.001) and helper (Pearson’s r = 0.64, FDR q-value < 0.001) T cells were enriched in the vicinity of tumor cells with higher GTT, whereas non-cycling helper (Pearson’s r = -0.97, FDR q-value < 0.001), cytotoxic (Pearson’s r = -0.92, FDR q-value = 0.011) and regulatory (Pearson’s r = -0.76, FDR q-value = 0.014) T cells were depleted, suggesting altered T cell dynamics and potential functional remodeling of the adaptive immune response. Within the macrophage compartment, SPP1-high (Pearson’s r = 0.87, FDR q-value = 0.009) macrophages became progressively more abundant with increasing GTT, whereas STAB1-high (Pearson’s r = -0.78, FDR q-value = 0.009) and CHIT1-high (Pearson’s r = -0.78, FDR q-value = 0.023) macrophages became less abundant.

**Figure 3:**
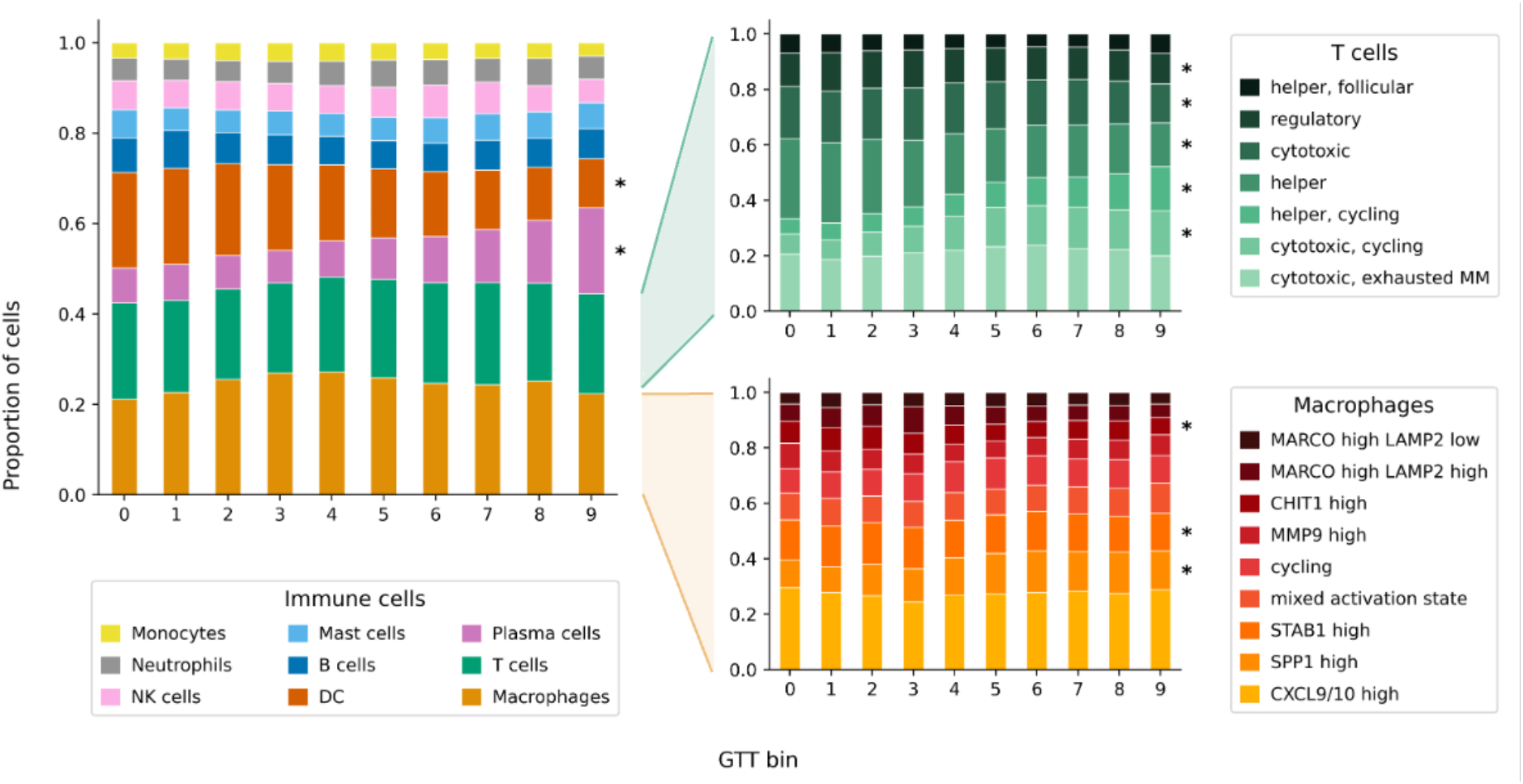
Immune cell composition in tumor microenvironment shifts with increasing GTT. Stacked bar plots show the relative proportions of the indicated immune cell subtypes within the tumor neighborhood (30 µm radius) across GTT bins (0–9). Left: Immune cell composition. Right: top: detailed T-cell subtype composition; bottom: detailed macrophage subtype composition. Asterisks (*) indicate statistically significant Pearson correlations after false discovery rate (FDR) correction (q ≤ 0.05), as determined by Pearson correlation analysis.

### Global Tumor Time of cancer cells is predictive of patient survival

Given that high-GTT tumor cells were associated with features of aggressive tumor biology, including increased proliferation and hypoxia, greater transcriptional/genomic complexity, altered cellular morphology and an immune-depleted microenvironment, we investigated whether the average GTT of a tumor was associated with patient prognosis.

GTT demonstrated consistent prognostic value for overall survival across the internal snRNA-seq cohort (C-index: 0.60), an independent external validation cohort comprising snRNA-seq data from 59 primary LUAD patients (C-index: 0.66), and the TCGA-LUAD bulk RNA-seq cohort (C-index: 0.61; Fig. 4A). Additionally, GTT retained robust discrimination even within stages I, II, and IV across all three cohorts. (Fig. 4B). For UICC stage III, GTT added similar prognostic information in the two external datasets but showed no additional risk stratification in the internal dataset.

**Figure 4:**
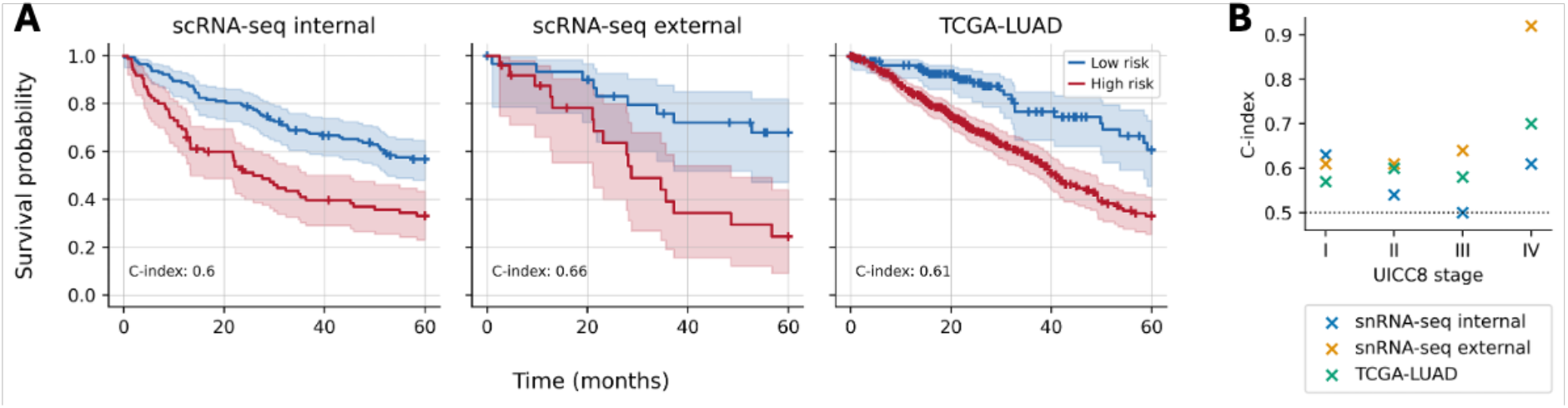
GTT stratifies patient survival across three independent datasets. (A) Kaplan-Meier survival curves for the internal snRNA-seq dataset (used for GTTimer fitting), external snRNA-seq dataset and TCGA-LUAD cohort (both unseen validation data). Patients were classified into low- and high-risk groups based on GTT. The optimal cutoff was determined using the log-rank test with permutation testing. Concordance index (C-index) for GTT is shown. (B) Concordance indices (c-indices) per dataset, stratified by UICC (8th edition) stage. The line at 0.5 indicates the expected performance of random risk stratification.

### Investigating GTT gene expression profiles with explainable AI

We used the explainable artificial intelligence (xAI) method layer-wise relevance propagation (LRP) to gain biological insights into how the expression of individual genes contributed to higher or lower GTT. To identify the major transcriptional programs underlying GTT, we applied non-negative matrix factorization (NMF) to the normalized absolute LRP scores, yielding two distinct gene programs which we annotated as early and late based on their activity across GTT (Fig. 5A, B). The early program contributed to lower GTT and was enriched for surfactant-associated and airway epithelial cell markers, including *SCGB3A2*, *SFTPD*, *SFTPC*, and *LAMP3*, consistent with an early tumor state in which LUAD cells retain transcriptional similarity to their cell of origin. The late program, contributing toward higher GTT, comprised genes associated with metabolic reprogramming, therapy resistance and poor outcome^24–27^. (Supplementary Fig. S2, Fig. 5C). Together, these findings suggest a transition from a differentiated epithelial state with similarity to normal epithelial airway cells toward increasingly advanced and aggressive tumor states with rising GTT.

Additionally, gene-level hazard ratios in TCGA-LUAD were significantly lower for early than for late program genes (Fig. 5D; p<0.001), indicating that early program genes were associated with lower risk.

**Figure 5:**
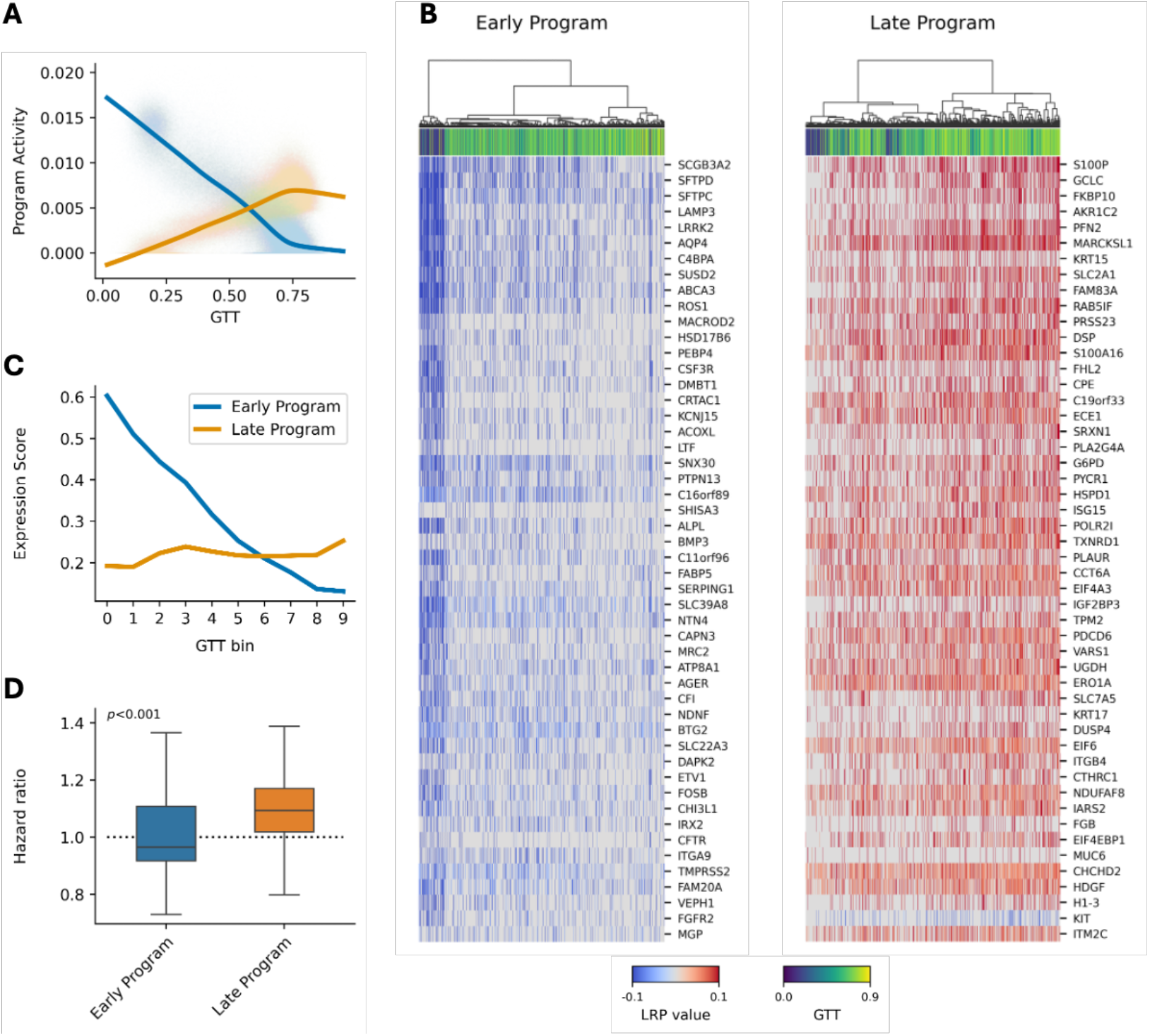
LRP identifies distinct gene programs defining GTT. (A) Activity of NMF-derived early and late gene programs across cells ordered by GTT. Solid lines indicate LOWESS-smoothed trends. (B) Heatmaps showing normalized LRP relevance scores for genes of the early (left) and late (right) programs across a random subset of 5,000 cells. Cells are ordered by hierarchical clustering (Ward linkage, Euclidean distance), and genes are ordered by mean absolute relevance (mean absolute normalized LRP across cells). Values were clipped to the 1st–99th percentile and transformed using a signed square-root transformation. Per program, only the top 50 of unique genes ranked by mean absolute relevance are shown. Column colors indicate GTT. (C) Mean expression scores of early and late programs across cells grouped by GTT bins. Scores were computed using Scanpy’s score_genes function based on log-normalized counts and averaged per GTT bin. (D) Gene-level hazard ratios in TCGA-LUAD, grouped by program. Differences between early and late gene programs were assessed using two-sided Welch’s t-test on log-transformed hazard ratios.

Notably, these LRP-defined programs showed no enrichment in ranked gene set enrichment analysis (GSEA) (Supplementary Tables S1, S2), suggesting that the underlying progression signals are not well represented by canonical pathway annotations.

To further characterize the biological processes associated with GTT, we performed GSEA on genes ranked by their correlation with GTT.

**Table 1:** Gene Set Enrichment Analysis (GSEA) results based on genes ranked by Spearman correlation coefficient with GTT. Gene sets are ordered by statistical significance (false discovery rate (FDR q-value) and nominal p-value). Enrichment score (ES) and normalized enrichment score (NES) are reported.

| **Pathway** | **ES** | **NES** | **p-value** | **FDR q-value** | **Gene overlap** |
|---|---|---|---|---|---|
| E2F Targets | 0.8277 | 3.0373 | < 0.001 | < 0.001 | 94/106 |
| G2-M Checkpoint | 0.8006 | 2.8987 | < 0.001 | < 0.001 | 85/96 |
| Myc Targets V1 | 0.7953 | 2.5403 | < 0.001 | < 0.001 | 44/45 |
| Mitotic Spindle | 0.6890 | 2.3281 | < 0.001 | < 0.001 | 40/60 |
| mTORC1 Signaling | 0.5400 | 1.9895 | < 0.001 | < 0.001 | 13/16 |
| Myc Targets V2 | 0.7785 | 1.9592 | < 0.001 | < 0.001 | 19/34 |
| Coagulation | -0.5305 | -2.0624 | < 0.001 | 0.0010 | 61/97 |
| Glycolysis | 0.4967 | 1.8080 | < 0.001 | 0.0012 | 67/104 |
| Unfolded Protein Response | 0.5854 | 1.7602 | < 0.001 | 0.0025 | 11/21 |
| Reactive Oxygen Species | 0.6316 | 1.7209 | 0.0025 | 0.0036 | 17/24 |

Genes increasing toward high GTT were strongly enriched for proliferative and cell cycle related programs, including E2F Targets (NES = 2.70, FDR q < 0.001), G2-M Checkpoint (NES = 2.66, FDR q < 0.001), MYC Targets V1 (NES = 2.44, FDR q < 0.001), and Mitotic Spindle (NES = 2.19, FDR q < 0.001). Enrichment of Unfolded Protein Response (NES = 0.65, FDR q < 0.001), Glycolysis (NES = 0.55, FDR q < 0.001), mTORC1 Signaling (NES = 0.54, FDR q < 0.001), Reactive Oxygen Species Pathway (NES = 0.70, FDR q < 0.001), and DNA Repair (NES = 0.67, FDR q < 0.001) further suggested increased metabolic activity, cellular stress responses, and genomic maintenance during tumor progression.

### Comparing GTTimer against benchmark methods

We compared GTTimer to two prior methods that estimate a common pseudotime across multiple samples, Phenopath and Monocle3 with prior batch correction, across datasets comprising tumor cells from randomly selected 2 (2,656 cells), 5 (6,880 cells), and 10 (16,059 cells) patients, as well as the full dataset with 227 patients (422,727 cells; Fig. 6A).

GTTimer outperformed the alternative methods in computation efficiency (Fig. 6A), requiring both less memory and substantially less runtime for the full cohort of 227 patients: For the full cohort, training GTTimer required 38.5 minutes, while inference required 4.6 seconds. In contrast, Monocle3 required 11.5 hours, indicating a substantially higher computational burden at scale. In terms of memory usage, GTTimer training and inference required 48 GB and 619 MB, respectively, whereas Monocle3 required 316 GB for the full cohort. Benchmarking tests for Phenopath for larger datasets than 10 patients were aborted after exceeding 50 hours of runtime.

Most, importantly, GTTimer demonstrated superior performance in survival prediction (Fig. 6B). Using the scores of Monocle3 to determine patient overall survival showed worse performance on the internal (C-index: 0.45) and external (C-index: 0.61) dataset than GTT (C-index = 0.60 and 0.66, respectively). These findings highlight that GTTimer combines robust scores with clinical relevance with computational scalability to very large single-cell datasets.

**Figure 6:**
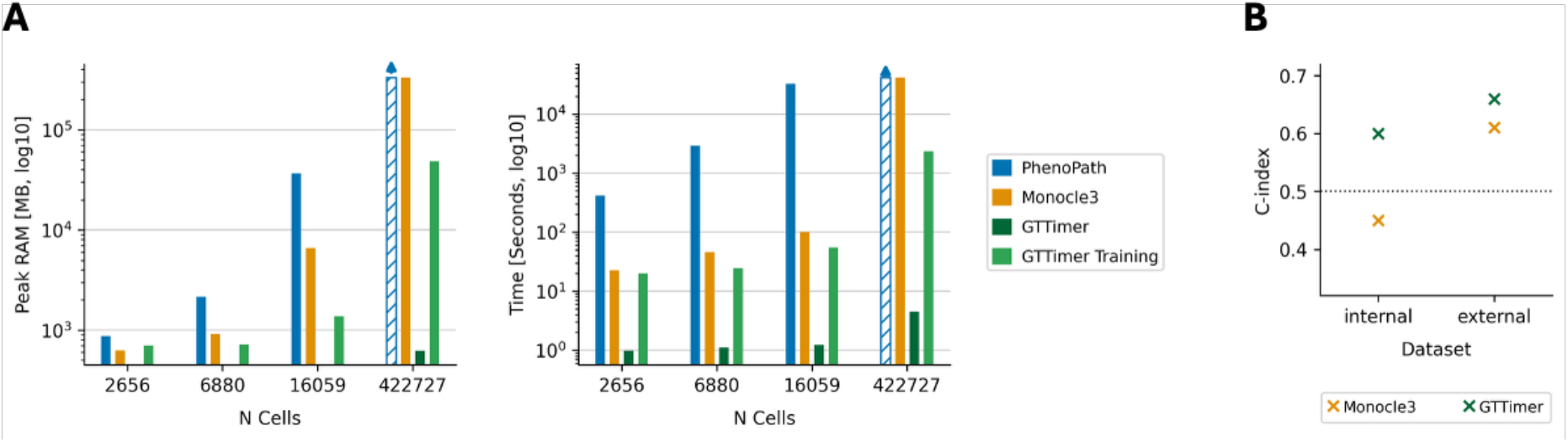
GTTimer outperforms existing approaches in scalability and predictive performance. (A) Peak RAM usage in MB (left) and runtime in seconds (right) for Phenopath, Monocle3 with batch-correction and GTTimer (training and inference), across datasets comprising 2, 5, 10, and 227 patients (up to 422,727 cells). Application of Phenopath to the full dataset was computationally infeasible, as indicated by the dashed bars with upward arrows. Y-axis is log scaled. (B) C-indices of Monocle3 and GTTimer for internal and external snRNA-seq datasets. Phenopath is not shown because it was not applicable to the full cohorts (see panel A). The line at 0.5 indicates the expected performance of random risk stratification.

## Discussion

In this study, we delineated the development of tumor cell states in LUAD by defining a shared measure of tumor progression across patients, overcoming the limitation of comparing pseudotime dynamics between samples in scRNA-seq datasets. By leveraging snRNA-seq data from 227 patients, we established GTTimer, a neural network-based approach enabling direct comparison of cellular states across samples, facilitating the study of tumor evolution at both intra- and inter-patient levels.

We showed that GTT is biologically and clinically meaningful and robust, aligning with established markers of tumor progression, such as advanced clinicopathological characteristics, proliferative and hypoxic signaling, enlarged cellular morphology, and immunologically depleted local microenvironments. GTT provides prognostic value for patient survival and outperforms existing methods in both computational efficiency and predictive accuracy, capturing progression-related signals beyond conventional clinical staging. This robustness extends across independent datasets, including both single-cell and bulk transcriptomic data, supporting the generalizability of GTTimer.

Previous efforts to delineate progressive changes in tumor cell states in LUAD have largely relied on Monocle3 with batch correction to align cells across patients^13–16^. In our analysis, such approaches yield progression measures with limited prognostic value, suggesting that batch correction alone does not fully resolve inter-patient variability and highlighting a limitation of conventional pseudotime methods in heterogeneous cohorts. A particular strength of GTTimer is its scalability to large datasets, outperforming benchmark approaches by far in terms of computational time and resources.

Our spatial analysis revealed reduced immune cell presence surrounding advanced tumor cells and a progressive shift in the composition of the immune microenvironment with increasing GTT. Previous spatial studies in LUAD have described a shift from adaptive to innate immune dominance during progression, characterized by depletion of cytotoxic T cells, NK cells, and B cells, and enrichment of macrophages and regulatory T cells^28–31^. While we similarly observed depletion of non-cycling cytotoxic T cells and expansion of SPP1-high macrophages, our analysis additionally revealed a decline in regulatory T cells and selective remodeling of macrophage subtypes, highlighting the increased resolution afforded by modeling tumor progression as a continuous process. In addition, dendritic cells decreased, whereas plasma cells increased with GTT, consistent with reports of plasma cell accumulation during LUAD progression^32,33^.

To dissect the molecular programs underlying GTT, we applied LRP to identify gene programs associated with early and late tumor states. The early program includes surfactant-associated genes (*SFTPD, SFTPC, LAMP3, ABCA3*)^34,35^, and regulators of homeostasis highly expressed in alveolar cells (*LRRK2*^36^*, AQP4*^37^). Many of these genes have been reported to exert tumor-suppressive functions (*SCGB3A2*^38^*, SUSD2*^39^*, HSD17B6*^40^*, LTF*^41^*, SNX30*^42^) or correlate with favorable clinical outcomes in LUAD (*C4BPA*^43^*, CRTAC1*^44^*, KCNJ15*^45^). In contrast, late program genes are associated with poor clinical outcome and promote key cancer hallmarks such as epithelial-mesenchymal transition (*MARCKSL1*^46^*, FAM83A*^47^), proliferative signaling (*FHL2*^48^*, ECE1*^49^*, G6PD*^50^), invasion and metastasis (*S100P*^24^*, PFN2*^51^*, FHL2*^48^*, PLA2G4A*^52^) and therapy resistance (*AKR1C2*^26^*, SLC2A1*^27^, DSP^53^). Importantly, the late program comprises *KRT15*^54^, a candidate for personalized and immunotherapeutic targeting, as well as *SLC2A1* (GLUT1)^55^, and *SLC16A3* (MCT4)^56^, emerging therapeutic targets involved in clinically actionable pathways such as glycolysis and lactate transport. Together, these findings delineate a progression from differentiated epithelial states toward aggressive tumor phenotypes, demonstrating that GTT captures biologically meaningful programs beyond canonical lineage markers.

Several limitations should be considered. While LRP provides interpretability, it does not establish causality^57^, and experimental validation will be required to confirm potential drivers of tumor progression. Moreover, although many genes associated with GTT were consistent with their reported roles in LUAD and other cancer types, the identified gene programs capture coordinated cellular states rather than the effects of individual genes. As a result, not every gene within the early or late program necessarily reflects the biological characteristics of that program. In addition, while the shared axis of malignant cell transformation enables direct comparison of cellular states across patients and tumors, it may not fully capture highly branched or parallel evolutionary trajectories.

Overall, our findings provide a single-cell resolution view of the malignant transformation of tumor cells in LUAD, revealing a coordinated transition from differentiated alveolar epithelial states to aggressive tumor phenotypes. The identified transcriptional programs highlight candidate regulators associated with distinct progression states. By enabling direct comparison of cellular states both across patients and within individual tumors, GTT provides a common framework for studying tumor evolution and intratumoral heterogeneity. Future studies may leverage similar approaches to investigate progression dynamics in other cancer types, helping to uncover conserved and tumor-specific mechanisms that shape disease progression. Ultimately, a deeper understanding of progression-associated cellular states and molecular programs may support patient stratification beyond conventional clinical staging and facilitate the identification of therapeutic vulnerabilities that emerge during tumor evolution.

## Methods

### Patients and Samples

We analyzed all available 227 LUAD cases from the discovery cohort of the previously generated LuCaS+ snRNA-seq atlas^20,21^. The LuCaS+ discovery cohort comprises clinically annotated FFPE tumor samples from NSCLC patients treated at two different German university hospitals, with snRNA-seq data generated using the 10x Genomics Flex protocol.

### snRNA-seq Data Processing

We leveraged a previously published snRNA-seq dataset generated by our group^20,21^. Filtering, normalization and celltype annotation were conducted as described in the original study. Batch correction across samples was performed using Harmony^58^ on PCA embeddings, computed from log-normalized expression values. UMAP was computed on the Harmony-corrected space using 50 principal components and 100 neighbors on a randomly subsampled set of 200,000 cells. DPT^22^ on tumor cells and type II pneumocytes was computed separately for each patient using the implementation provided in Scanpy (v1.11.0)^59^. Type II pneumocytes were included to anchor the root in the presumed cell of origin and orient pseudotime from non-malignant epithelial cells toward malignant cells. Root cells were defined using the average expression profile of non-malignant epithelial cells. After DPT computation, only tumor cells with finite DPT were retained for all downstream analysis. Cell cycle phase was assigned using the score_genes_cell_cycle() function in Scanpy with S-phase and G2/M-phase gene sets obtained from the Seurat package (v5.3.1)^60^. Hypoxia scores were computed based on the Tardon^61^ hypoxia gene signature by calculating the mean expression of the signature genes using the R package sigscores (v0.0.0.9000)^62^. CytoTRACE2 (v1.1.0)^63^ was applied to infer a potency score for each cell, reflecting its predicted developmental potential. Copy number variation (CNV) inference was performed by applying inferCNV (v1.23.0)^64^ to raw count matrices for each patient independently, using non-malignant epithelial, stromal and immune cells as the normal reference. The analysis was performed without HMM-based state prediction, and default smoothing and denoising parameters were used. CNV score per cell was defined as the fraction of genes whose expression deviates from the reference by more than ±5%.

### Linear Mixed Effects Model

For each gene, DPT was modeled as a function of gene expression using a linear mixed-effects model to account for inter-patient heterogeneity:

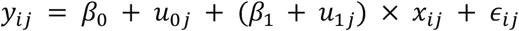

where:

- *y_i_*_j_: DPT of cell i in patient j

- *x_i_*_j_: gene expression of cell i in patient j

- *β*_0_, *β*_1_ : global intercept and expression coefficient (fixed effect)

- *u*_0j_, *u*_1j_: patient-specific intercept and slope (random effect)

- *ε_i_*_j_: residual error.

Genes were ranked by model performance, defined by the square root of the mean squared residuals (RMSE), and the top 10% of genes were selected for further analysis.

### Neural Network Model for GTT Prediction

We implemented a neural network in PyTorch (v2.6.0)^65^ to estimate GTT from gene expression data. During training, the model was optimized in a supervised manner using per-cell DPT values as the target. Gene expression served as input to a multi-layer perceptron (MLP) consisting of a sequential of a fully connected linear layer, a layer normalization, a GELU activation, followed by a second fully connected layer that produced a latent progression score. Dropouts were applied at the input layer and after GELU. To account for inter-patient variability in DPT scales, the patient ID per cell was mapped to a trainable embedding (using torch.nn.Embedding) containing two patient-specific parameters representing intercept and slope. During training, these embedding parameters were combined with the latent progression score in the final model layer to reconstruct the observed DPT according to:

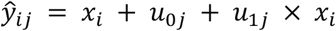

where:

- *ŷ*,*_i_*_j_: reconstructed DPT of cell i in patient j

- *x_i_*: latent progression score of cell i, predicted by the MLP

- *u*_0j_, *u*_1j_: patient-specific intercept and slope for patient j, learned via the embedding layer.

GTT was obtained directly from the MLP (*x_i_*) by omitting the patient-specific embedding layer, thereby removing the patient-specific scaling and offset learned during training.

### Model Training and Hyperparameter Optimization

The dataset was randomly split at the patient level into a training set (n = 192 patients) and a validation set (n = 35 patients). Model parameters were optimized using the Adam optimizer^66^ by minimizing the Smooth L1 loss between reconstructed and reference DPT values together with an L2 regularization term on the patient-specific embedding parameters (intercepts and slopes). The contribution of the regularization term was controlled by the hyperparameter *λ_RE_*, which penalized large patient-specific effects and thereby encouraged the latent progression score to capture progression shared across patients. Training was performed for a maximum of 25 epochs, with early stopping based on the validation objective using a patience of 5 epochs.

Hyperparameter tuning was performed using Ray Tune^67^ with a grid search over the learning rate, weight decay, batch size, dropout rate, number of hidden units, gradient clipping norm, and the regularization parameter *λ_RE_*. Hyperparameter configurations were evaluated on the validation set using a validation objective defined as the sum of the Smooth L1 loss and one minus the mean per-patient Spearman correlation between reconstructed and reference DPT values. This validation objective balanced accurate reconstruction of DPT with preservation of the patient-specific ordering of cells. The trained model achieving the lowest validation objective was retained and used directly for all downstream analyses.

The selected hyperparameter configuration comprised a learning rate of 0.001, weight decay of 0.001, a batch size of 2,500 cells, 128 hidden units, a dropout rate of 0.4, gradient clipping with a maximum norm of 5, and *λ_RE_*=0.01. Using these hyperparameters, the final model was retrained for 25 epochs.

### Spatial Transcriptomics

Spatial transcriptomics was performed on tissue microarrays (TMAs) comprising 651,763 tumor cells of 152 matched LUAD cases. For each patient, two 1.5 mm tumor cores were included. Xenium in situ analysis was performed using the 10x Genomics Xenium Human Lung Panel supplemented with a custom add-on panel of 100 genes selected to capture cellular pathway activity and tumor-microenvironment interactions. Following Xenium analysis, the same tissue sections were H&E stained to enable direct comparison of spatial transcriptomic data with histomorphology. After integration of the spatial dataset and snRNAseq data of matched samples with SpaGE^68^, GTT was projected onto the Xenium tumor cells by the mean of 20 nearest neighbors, weighted by distance.

### Clinical Endpoints

Overall survival was defined as the duration from tumor diagnosis to death from any cause. Patients without a recorded death were censored at the time of their last follow-up. Follow-up was administratively censored at 60 months, such that patients remaining at risk beyond 60 months were censored at that time.

Prognostic performance was quantified using the concordance index (C-index) estimated from a Cox proportional hazards model. For visualization of survival differences, patients were stratified into low- and high-risk groups using the optimal cutoff determined by permutation testing, and Kaplan-Meier survival curves were generated accordingly.

### External Validation Data

To assess external validity, we applied the trained GTTimer to an independent snRNA-seq dataset comprising 50 patients previously generated and published by our group^20,21^, as well as to TPM-normalized expression data from the TCGA-LUAD bulk transcriptomic cohort, including 501 patients with available clinical follow-up. The external snRNA-seq dataset had undergone preprocessing and quality control as described in the original publication and was additionally used to validate the linear mixed-effects models. DPT and linear mixed-effects modeling were computed in this cohort using the same analytical pipeline and parameter settings as applied to the main dataset.

### Explaining GTTimer

We used Layer-wise Relevance Propagation (LRP)^69^ for explaining GTTimer. LRP is a post hoc explanation method that propagates model prediction layer by layer to the input. By doing so, each input feature is assigned a scalar value, quantifying its contribution to the model prediction. The layer-wise propagation follows so-called LRP-rules^70^.

LRP follows a general relevance conservation rule, i.e., is 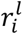 is the relevance of the i-th neuron in layer l, 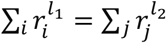 for any two layers *l*_1_ and *l*_2_. Additionally, LRP relevance propagation from layer (l+1) to layer l has a general form of:

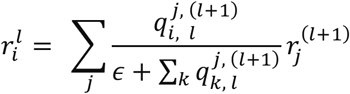

where 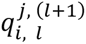 is the contribution of neuron i of layer l to neuron j of layer (l+1) and ɛ is a small value for stabilizing computations.

For linear layers, we used LRP-γ rule^70,71^, where 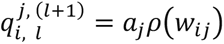, with *a*_j_ being the activation of j-th neuron in layer l, *w_i_*_j_ is the weight of connecting neuron i of layer l to neuron j of layer (l+1) and 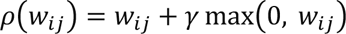, *w_i_*_j_7. In our experiments we used γ=0.

For LayerNorm layer, we use LN-rule^72^, which includes detaching the standard deviation in the denominator of the normalization term.

For the Gelu layer, we followed the MambaLRP framework^73^, which includes detaching the cumulative distribution function of the Gaussian distribution in the activation function.

LRP scores were normalized per cell by dividing by the sum of absolute relevance values across all genes, ensuring comparability across cells. To identify latent gene programs underlying model predictions, we applied non-negative matrix factorization (NMF) to the absolute normalized LRP scores. NMF was performed with two components using Nonnegative Double Singular Value Decomposition (NNDSVD) initialization and a maximum of 500 iterations, resulting in a decomposition into program activity per cell (W) and gene weights per program (H). For each program, genes were ranked according to their corresponding weights in H, and preranked GSEA was performed using all genes as the ranked background. In parallel, the minimal subset of top-ranked genes explaining at least 70% of the cumulative program weight was selected. Genes appearing in both programs were subsequently excluded, retaining only genes unique to each program for visualization and interpretation. Gene programs were annotated as early and late according to their activity patterns across GTT.

### Gene Set Enrichment Analysis

Pre-ranked GSEA was performed using the MSigDB Hallmark gene sets on three ranked gene lists. The first and second ranked gene lists consisted of genes ranked by their absolute NMF weights in the early and late programs, respectively. The third ranked gene list consisted of genes ranked by their Spearman correlation coefficient between log-normalized gene expression and GTT. For all three ranked gene lists, GSEA was performed using default parameters, restricting gene sets to a minimum size of 5 genes and a maximum size of 500 genes.

### Benchmarking

To assess computational performance and scalability, we benchmarked our method against Phenopath^17^ and Monocle3^12^ with batch correction^74^. All methods were applied to the same random subsets (2, 5, 10 patients) and the full cohort of the internal snRNA-seq dataset, restricted to tumor cells and including all genes. DPT computation for GTTimer additionally included pneumocytes type II, to anchor the root in the presumed cell of origin, consistent with the primary analysis. For Monocle3 and GTTimer, root cells were defined using the average expression profile of non-malignant epithelial cells.

For Monocle3, measurements included normalization, batch correction with align_cds(), dimensionality reduction, clustering, principal graph learning, and cell ordering. For GTTimer, training benchmarks included computation of DPT and training of the final model using the best hyperparameter configuration.

Computational performance was evaluated in terms of peak memory usage (MB) and runtime (seconds). All methods were executed on a machine with AMD EPYC 75F3 32-Core Processor, 2TB RAM, to ensure comparability.

Application of Phenopath to the full dataset was computationally infeasible. Analyses of datasets comprising more than 10 patients were terminated after exceeding 50 hours of runtime. Consequently, Phenopath was not evaluated on the full dataset.

### Statistical Analysis

All reported p-values for Pearson and Spearman correlation analyses were two-sided, and statistical significance was defined as p < 0.05. For analyses involving multiple comparisons, false discovery rate correction was applied using the Benjamini-Hochberg method. Two-sided Welch’s t-test was used to compare the log-transformed hazard ratios of early and late program genes in the TCGA-LUAD cohort.

## Contributions

Conceptualization: PK

Methodology: DK, PJ, PK, MJI

Formal analysis: DK

Investigation: all authors.

Resources: PJ, FK, KRM

Data curation: DK, PJ, FES

Writing - original draft: DK, PJ, PK

Writing - review and editing: all authors.

Visualization: DK

Supervision: PJ, PK

## Data Availability Statement

The internal and external snRNA-seq datasets analyzed in this study are available in Zenodo with the identifier doi:10.5281/zenodo.21877226. The supporting Xenium dataset is not publicly available due to data-sharing restrictions.

## Code Availability Statement

The custom code used in this study will be made publicly available upon publication.

