## Supplementary Material for "Learning the Malignant Transformation of Lung Adenocarcinoma Cells Using Explainable AI"

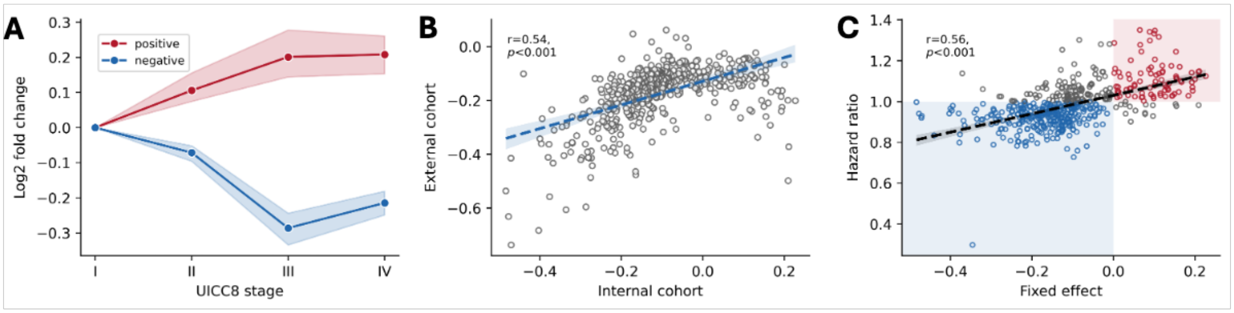


**Figure S1**: **Gene effects estimated from linear mixed effects models.** (A) Log2 fold expression changes across UICC8 tumor stages relative to stage I. Genes with positive fixed effect (red) show increasing expression, whereas genes with negative fixed effect (blue) exhibit reduction. Shaded areas indicate 95% CI. (B) Fixed effects in the internal cohort correlate with those in the external snRNAseq cohort (Pearson's $r = 0.54$, $p < 0.001$), based on independently fitted mixed effects models. (C) Fixed effects correlate with hazard ratios in the TCGA-LUAD cohort (Pearson's $r = 0.56$, $p < 0.001$; Fig. 1C), indicating a prognostic relevance. Point colors indicate the direction of the fixed effect (blue: negative, red: positive), and background shading reflects hazard ratio direction (blue: HR < 1, red: HR > 1). For all analysis, top 10% of genes (n=500) with best model fit were used.

| Pathway | ES | NES | p-value | FDR q-value | Gene overlap |
|---|---|---|---|---|---|
| Inflammatory Response | 0.7411 | 1.2480 | 0.0010 | 0.2686 | 26/91 |
| IL-6/JAK/STAT3 Signaling | 0.7588 | 1.2646 | 0.0018 | 0.3056 | 21/66 |
| Apical Surface | 0.7294 | 1.1917 | 0.0626 | 0.3319 | 19/34 |
| Apoptosis | 0.7215 | 1.2165 | 0.0030 | 0.3340 | 33/97 |
| Reactive Oxygen Species Pathway | 0.6959 | 1.1722 | 0.0114 | 0.3381 | 51/104 |
| UV Response Dn | 0.7012 | 1.1788 | 0.0146 | 0.3488 | 35/86 |
| Protein Secretion | 0.6653 | 1.1207 | 0.0654 | 0.3656 | 30/101 |
| Hedgehog Signaling | 0.6653 | 1.1232 | 0.0542 | 0.3750 | 28/105 |
| p53 Pathway | 0.8033 | 1.2925 | 0.0218 | 0.3787 | 9/21 |
| Androgen Response | 0.6725 | 1.1308 | 0.0496 | 0.3874 | 44/95 |

**Table S1:** GSEA results based on genes ranked by their contribution to the early program. Gene sets are ordered by statistical significance (false discovery rate (FDR q-value) and nominal p-value). Enrichment score (ES) and normalized enrichment score (NES) are reported.

| Pathway | ES | NES | p-value | FDR q-value | Gene overlap |
|---|---|---|---|---|---|
| TNF-alpha Signaling via NF-kB | 0.7206 | 1.0605 | 0.1674 | 0.7484 | 59/144 |
| Peroxisome | 0.6994 | 1.0106 | 0.4596 | 0.7581 | 20/36 |
| TGF-beta Signaling | 0.7092 | 1.0137 | 0.4617 | 0.7610 | 13/21 |
| KRAS Signaling Up | 0.6820 | 1.0035 | 0.4822 | 0.7840 | 30/146 |
| Pancreas Beta Cells | 0.7119 | 1.0141 | 0.4839 | 0.7847 | 5/14 |
| Bile Acid Metabolism | 0.7329 | 1.0608 | 0.2822 | 0.7920 | 12/41 |
| Cholesterol Homeostasis | 0.7266 | 1.0517 | 0.3058 | 0.7944 | 15/44 |
| Estrogen Response Early | 0.6911 | 1.0157 | 0.4050 | 0.8009 | 44/126 |
| DNA Repair | 0.7148 | 1.0260 | 0.4282 | 0.8099 | 4/24 |
| Estrogen Response Late | 0.7231 | 1.0634 | 0.1470 | 0.8161 | 40/136 |

**Table S2:** GSEA results based on genes ranked by their contribution to the late program. Gene sets are ordered by statistical significance (false discovery rate (FDR q-value) and nominal p-value). Enrichment score (ES) and normalized enrichment score (NES) are reported.

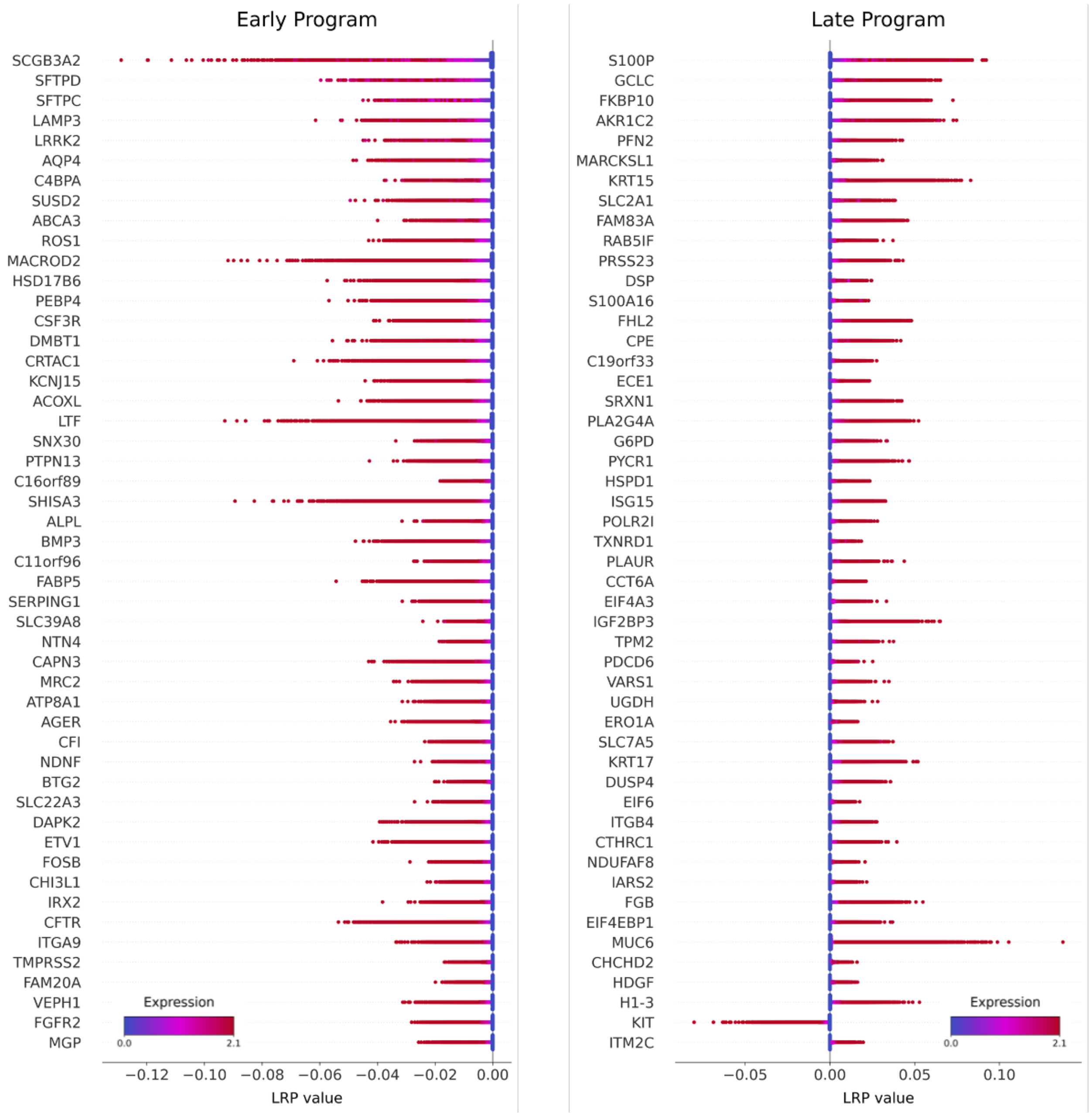


**Figure S2: LRP contributions are primarily driven by elevated gene expression.** Relationship between gene expression and LRP contribution of early program genes (left) and late program genes (right). Per program, top 50 of unique genes ranked by mean absolute relevance are shown. Low expression levels were associated with near-zero LRP values, whereas high expression increased absolute LRP contributions.
